# Proofreading Is Too Noisy For Effective Ligand Discrimination

**DOI:** 10.1101/2023.01.13.523988

**Authors:** Duncan Kirby, Anton Zilman

**Author notes:** A.Z. and D.K. devised the project. D.K. performed the research. A.Z. and D.K. wrote the paper. The authors declare no competing interests.

## Abstract

Kinetic proofreading (KPR) has been used as a paradigmatic explanation for the high specificity of important biological processes including ligand discrimination by cellular receptors. Kinetic proofreading enhances the difference in the mean receptor occupancy between different ligands, thus potentially enabling better discrimination. On the other hand, proofreading also attenuates the signal, increasing the relative magnitude of noise in the downstream signal. This can interfere with reliable ligand discrimination. To understand the effect of noise on ligand discrimination beyond the comparison of the mean signals, we formulate the task of ligand discrimination as a problem of statistical estimation of the molecular affinity of ligands. Our analysis reveals that proofreading typically worsens ligand resolution which decreases with the number of proofreading steps under most commonly considered conditions. This contrasts with the usual notion that kinetic proofreading universally improves ligand discrimination with additional proofreading steps. Our results are consistent across a variety of different proofreading schemes, suggesting that they are inherent to the KPR mechanism itself rather than any particular model of molecular noise. Based on our results, we suggest alternative roles for kinetic proofreading schemes such as multiplexing and combinatorial encoding in multi-ligand/multi-output pathways.

## 1. Introduction

Cells send and receive signals in the form of molecular ligands that bind corresponding cell surface receptors. Ligand-receptor binding activates intracellular cascades of molecular modifications that eventually lead to cellular response. The ability of the cell to respond distinctly to different ligands depends on the ability of the receptor to discriminate between different ligands. In the simplest — and very common — scenario, discrimination relies on the differences in the receptor binding affinities between different ligands: cognate ligands bind the receptor more strongly (and for a longer time) than non-cognate ligands. At equilibrium, the ratio of the receptor occupancies by different ligands present at the same concentration is expected to be proportional to 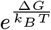 where Δ*G* is the difference in binding free energy between the ligands (1–3). However, the fidelity of many important biological processes has been observed to be higher than what can be expected from the measured differences in the equilibrium binding free energies between ligands (1, 4–9). In one prominent examples, T cells appear to be able to distinguish between self-peptides and foreign antigens which may differ by as little as a single amino acid mutation (as little as a two-fold difference in affinity) (3, 4, 10–14). Such extreme specificity of the cellular response to cognate ligands is frequently explained in terms of a mechanism known as kinetic proofreading (KPR) (3, 7, 13–16).

The KPR mechanism enhances occupancy differences between ligands beyond the equilibrium factor 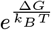 via introduction of one or more intermediate bound states between the initial binding and the final state where the output signal is produced (see Fig. 1A). The delay introduced by these additional states increases the probability of the ligand to dissociate from the receptor before the signal is produced, so that that the weakly binding ligands rarely stay bound long enough to reach the signaling state and produce an output. Magnifying the differences between the ligands comes at the expense of energy consumption (typically in the form of ATP hydrolysis) to ensure irreversible unbinding of the ligand from the proofreading states of the receptor (3, 16–18).

Kinetic proofreading was originally proposed by Hopfield and Ninio as a mechanism to explain the fidelity of DNA replication and protein biosynthesis and is believed to describe the proofreading step of the endonuclease and tRNA proofreading by the ribosome (1, 2, 19). McKeithan subsequently adapted multi-step KPR as an explanatory mechanism behind the remarkable specificity of the T cell receptor (TCR) in recognizing foreign versus self antigens (3). Subsequently, proofreading models and their modifications have been applied to explain additional features of the T cell response such as the speed and antagonism, while other works explored energetic and time constraints of generalized proofreading schemes (15–17, 20–23). KPR models have also been applied to other receptor signaling systems including the Fc*E*RI-IgE system, RecA assembly, and various intracellular signal cascades (7, 8, 13, 14, 16, 18, 24–31). This body of work established the common view of the generalized sensing problem: discrimination of two different ligand types (such as selfvs. non-self peptides for T cells) based on a signal produced by the ligand binding to the receptor.

In the context of cellular sensing, receptor resolution for different ligands is commonly measured by the ratio of the corresponding average occupancy probabilities of the signaling state of the receptor, or by the ratio of physically related quantities such as the average time spent in the signaling state, or the average amount of the produced downstream signaling output (1–3, 13, 21). In this paper, we will refer to this metric as the Hopfield-Ninio (HN) metric.

The HN metric characterizes the resolution in terms of ensemble average properties, and thus does not take into account the stochasticity in the molecular processes of the ligand binding and unbinding, proofreading steps, and the subsequent signaling reactions. However, as known in the context of concentration sensing, this stochasticity can have substantial effects on the sensing accuracy (32–39). As illustrated in Fig. 1B, the ability of a cell to resolve two different ligands depends not only on the separation of the mean outputs of the receptor, ⟨*n*_*i*_ ), but also on the overlap of the probability distributions of their outputs, *P* (*n* |*k*_off,i_), where ligand types are indexed by *i*, and *k*_off,i_ is the ligand-receptor unbinding rate which uniquely determines the ligand affinity. The problem of resolving two ligands is conceptually analogous to the two-point resolution problem familiar in optics, with the point spread function of each light source corresponding to the distribution *P* (*n* |*k*_off,i_) in molecular signaling (40, 41). As we show below, the effects of noise for KPR receptors can be especially acute because enhancement of the differences in mean signal between the ligands is accompanied by attenuated receptor output, which amplifies the role of noise (3, 37, 42). This signal attenuation is an inherent property of the KPR mechanism, and KPR is most commonly invoked as a mechanism of discrimination between a cognate ligand (weakly attenuated) and a non-cognate ligand (strongly attenuated) (43).

**Fig. 1.**
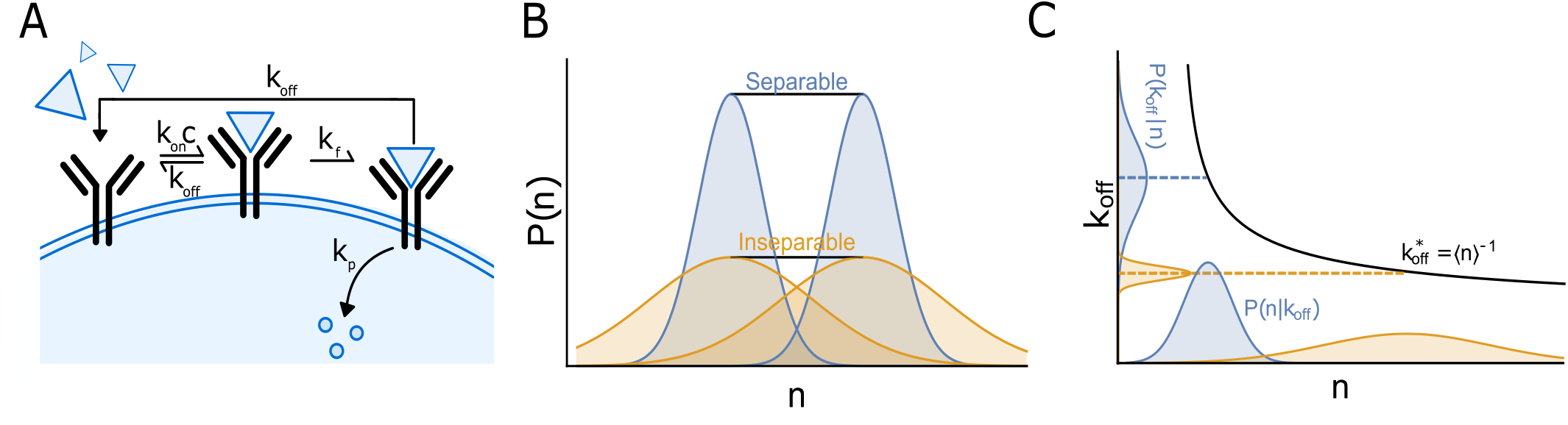
Ligand discrimination performance depends on molecular noise. A) Schematic diagram of a 1-step KPR receptor. The receptor in the unbound state can bind a ligand at a rate *k*_on_*c*, where *c* is the ligand concentration; the ligand dissociates from a bound receptor at a rate *k*_off_. In the bound state, the receptor may transition to the signaling state at a rate *k*_*f*_, from which it can return to the unbound state again at rate *k*_off_. While in the signaling state the receptor produces output molecules at a rate *k*_*p*_. B) Ligand discrimination depends on the separability of output distributions dictated by the distance between their means and their widths. The blue distributions are easily separable, while the orange distributions that have the same distance between their means are inseparable due to greater variance in *n*. C) Ligand discrimination can be formulated as the estimation of the dissociation rate *k*_off_ from the output *n*. The noise in the output *n* described by the distribution *P* (*n*|*k*_off_) translates into a distribution of errors in the estimate for *k*_off_ which is centered on 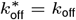. We plot the likelihood, *P* (*k*_off_|*n*), of *k*_off_ for a particular *n*. The estimation function (black curve) determines how variance in *n* translates into variance in 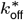; we demonstrate two regimes (orange vs. blue) where the variance in *n* has less or more effect on the variance in 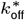 respectively.

It is important to distinguish the effect of the intrinsic molecular noise on signal discrimination from the role of the potential extrinsic constraints such as the energy consumption rate or the speed of the cellular decision-making. Extrinsic constraints represent a trade-off, with better resolution available if only the costs can be met or else sacrifice the resolving power (39, 44–46).

By contrast, as we show in this paper, the intrinsic and unavoidable molecular noise of proofreading directly affects the ligand resolution capability of the receptor. In general, the maximal information about the nature of a ligand is encoded in the whole sequence of binding and unbinding events during the measurement time. However, information processing in cellular signaling is constrained by molecularly realizable mechanisms which by necessity reduce the information available to the cell. In this paper we specifically focus on the effect of proofreading on specificity and ligand resolution and compare the resolution of a proofreading receptor to a non-proofreading receptor using mathematical models informed by biologically realistic signaling modalities.

The role of molecular noise in receptor sensing has been investigated in a variety of contexts. Extending the pioneering work by Berg and Purcell, (32, 34–36, 47–53) adopted the Fisher Information and the associated framework of statistical estimation theory to quantify how receptor occupancy fluctuations fundamentally limit the precision of estimating ligand concentration. Other works have investigated the specificity of response to a particular ligand using information theoretic tools (18, 22, 36, 48, 54–59). Selecting between these various approaches to account for noise in ligand processing performance depends on the question at hand.

In this paper, we use the mathematical framework of statistical estimation theory to account for the role of molecular noise in quantifying ligand specificity and resolution (53, 60). We view specificity as the precision of estimating the ligand binding affinity to the receptor. Accordingly, we treat the ligand discrimination task as a problem of separating the affinity estimates for different ligands, and we construct the appropriate quantitative resolution metric. The metric goes beyond a simple comparison of the mean receptor output and describes precisely the effect of intrinsic proofreading noise on the ligand discrimination task.

Our results reveal that once the intrinsic molecular noise is accounted for, the proofreading receptor’s resolution ability is lower than that of a non-proofreading receptor in most of the parameter space, including the the regime of low ligand concentration and strong proofreading where KPR is commonly invoked. We show that proofreading improves resolution only in a narrow non-classical area of the parameter space with a low number of proofreading steps and weak proofreading strength. As proofreading steps are added, the resolution decreases relative to a non-proofreading receptor because KPR amplifies the molecular noise more strongly than it separates the means of the receptor outputs for different ligands. This effect raises new questions about the value of KPR as a mechanism for unconstrained enhancement of ligand discrimination performance, and suggests that molecular noise limits the feasibility of KPR applications which rely on multiple proofreading steps to achieve the desired specificity (18, 21, 27, 61). Despite limited number of quantitative experimental measurements of the KPR parameters, recent results showing that the effective number of the proofreading steps for the TCR is relatively small (13, 14) and the proofreading strength is moderate (14) are consistent with our results.

The paper is structured as follows. In Section 2A we briefly review the KPR model. In Section 2B we show how the intrinsic molecular noise affects the output distribution in the KPR model and examine the signal-to-noise ratio of the model. In Section 2C we formulate the problem of specificity as the statistical estimation of ligand affinity and examine how receptor occupancy fluctuations affect the ability to identify ligands, an important precursor for considering the problem of ligand discrimination. In Section 2D we introduce a metric for ligand resolution derived from statistical estimation theory. In Section 2E we show that resolution enhancement by KPR is severely restricted due to the amplification of the relative noise by the KPR scheme. In Section 2F we investigate alternative KPR schemes and metrics to show that our results are inherent to the proofreading mechanism itself rather than a particular choice of the noise model of a discrimination performance metric. We conclude with a discussion of our results in the context of previous work, and their consequences for the understanding of the roles of KPR in cellular signaling.

## 2. Results

### A. The KPR Model

We model a KPR receptor as a system with *N* +2 states. These states represent the unbound receptor (state 0), the bound receptor in each of the *N* proofreading states, and the final signaling state of the receptor (numbered as *N* + 1, see Fig. 1A). Thus, *N* denotes the number of proofreading states and *N* = 0 corresponds to a non-proofreading receptor.

The ligand, present at concentration *c*, can bind an unoccupied receptor with the overall rate *k*_on_*c*. Once bound by ligand, the receptor can sequentially transition from the initial bound state to each subsequent proofreading state at a rate *k*_*f*_. The ligand can unbind from the receptor in any state at a rate *k*_off_. We assume that the ligand-receptor association rate *k*_on_ is diffusion limited and is independent of the ligand identity so that only the unbinding rate is related to the ligand-specific binding free energy (i.e. 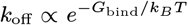). Note that the equilibrium dissociation constant, also known as the affinity, is *K*_*d*_ = *k*_off_*/k*_on_ and in our case is uniquely determined by *k*_off_. In the terminal bound state the receptor produces downstream signaling molecules at a rate *k*_*p*_, accumulating *n* output molecules over time *t*. In the following, we measure time in units of 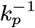 throughout this paper. This type of receptor output model is a common representation of a signaling mechanism by phosphorylation of intracellular molecules by a receptor-bound kinase (62–65). Other choices of receptor output models are investigated in Section 2F.

Ligand binding and unbinding are subject to intrinsic thermal and chemical fluctuations so that the transitions between the states in the model are probabilistic with the average rates described above. Together with the stochastic fluctuations in the production of the output molecules, this intrinsic noise leads to the distribution *P* (*n* |*k*_off_) of possible outputs, *n*, from the receptor in response to a ligand with the unbinding rate *k*_off_ (see Section 2F for models with minimal chemical noise). At long times — defined as min(*k*_off_*t, k*_*f*_ *t*) ≫1 — or fast phosphorylation (*k*_*p*_*t ≫*1) the output distribution for an *N* - step receptor is well approximated by the Normal distribution 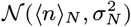 with the mean output from the receptor ⟨*n)*_*N*_ and its variance 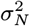 (53). In some situations, the cell needs to make a ligand discrimination decision in a short time, before the output distribution has reached the Normal distribution. This regime is analysed in Section 2F and the main results of the paper are recapitulated in this regime as well.

The expressions for the mean and the variance can be derived from the Master equation of the KPR model (see Materials and Methods):

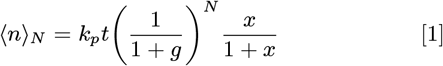

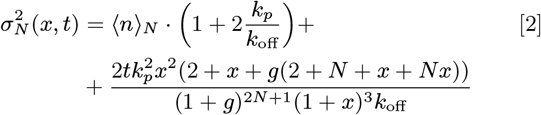

where *N* is the number of proofreading steps (*N* = 0 is a non-proofreading receptor), *g* = *k*_off_*/k*_*f*_ is the proofreading strength, and *x* = *c/K*_*d*_ is the non-dimensionalized ligand concentration. Equation [1] expresses the fact that at long times the number of output molecules produced by time *t* is simply the product of *k*_*p*_*t* and the probability to be in the signaling state *P*_*N* +1_ = *x/* (1 + *x*)(1 + *g*)^*N*^. Equation [2] reflects the fact that the variance in the number of output molecules is the sum of the intrinsic variance of the receptor occupancy distribution and the variance of the Poissonian process producing output molecules at rate *k*_*p*_ (53).

### B. Noise in the receptor output space

As illustrated in Fig. 1B-C, the width of the distribution *P* (*n k*_off_) is an important factor controlling the ability to resolve two ligands. In this section we explore how proofreading affects the width of the receptor output distribution.

The signal-to-noise ratio (SNR), 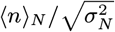, is commonly used to quantify the effect of noise in any signaling system, and is higher when the signal distribution is more tightly centered about its mean. In Fig. 2A we show the SNR for a KPR receptor as a function of the number of proofreading steps, *N*, and the proofreading strength, *g* = *k*_off_*/k*_*f*_. Critically, the SNR *decreases* with increasing proofreading strength (larger *k*_off_ or smaller *k*_*f*_). This trend can be understood from Eqs. 1-2, which show that in the high *g* limit the mean signal, ⟨*n)* _*N*_, scales like 𝒪 (1*/g*) while the noise, 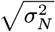, scales like 𝒪 (1*/g*^*N/*2^). Thus, the SNR scales as 𝒪 (*g*^−*N/*2^), and strong proofreading yields a lower SNR. Physically, the decrease in the SNR with increasing proofreading strength stems from the fact that a higher proofreading strength increases the variance in both the receptor occupancy distribution and the Poissonian process producing output molecules (see Eq. 2), amplifying the relative noise strength. We show in Section 2F that the increase in the receptor occupancy variance is inherent to KPR and sufficient to decrease the receptor’s signal processing performance.

**Fig. 2.**
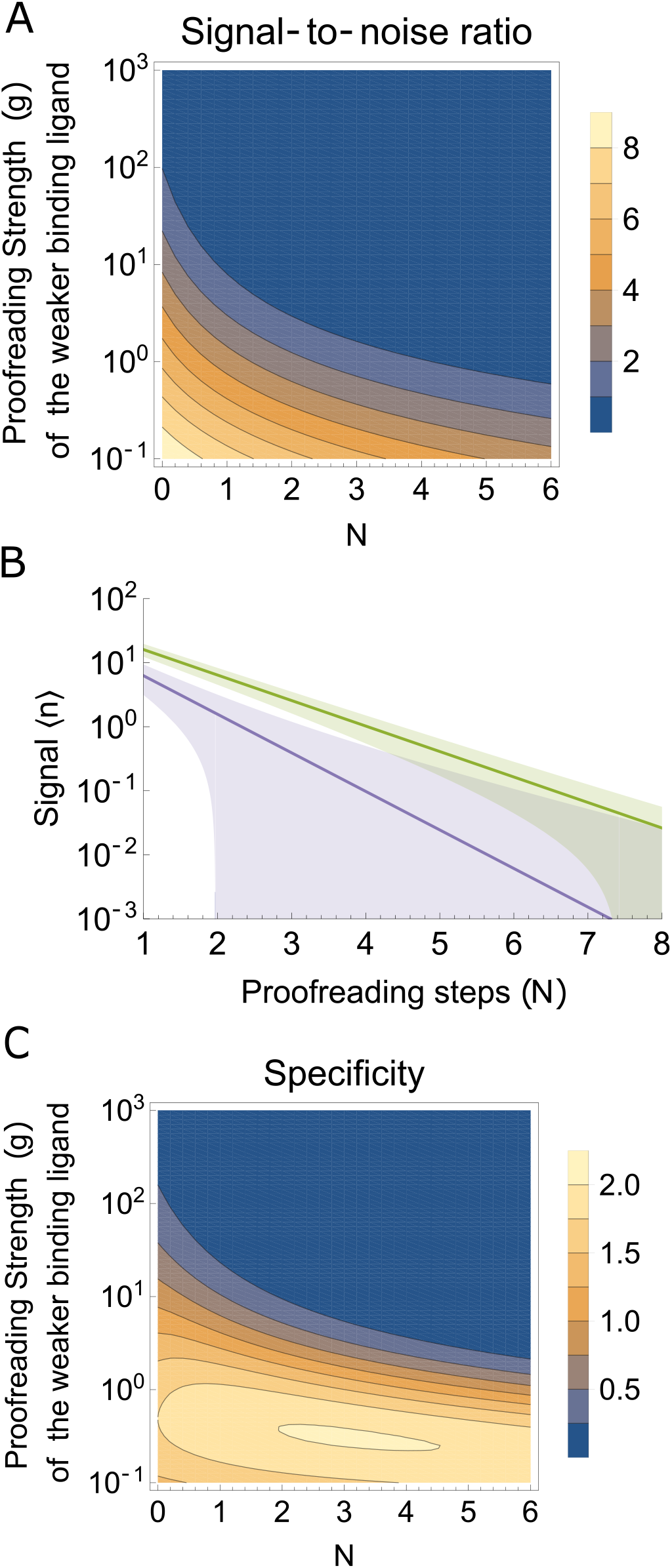
Proofreading does not improve the SNR or specificity. A) The signal-to-noise ratio ⟨*n /σ*⟩ _*n*_ for a KPR receptor as a function of the number of proofreading steps *N* and proofreading strength *g* = *k*_off, 1_*/k*_*f*_, with *k*_on_*c/k*_*p*_ = 1, *k*_*f*_ */k*_*p*_ = 1, *k*_*p*_*t* = 100, and *k*_off,2_ = 0.1*k*_off, 1_. B) The noisy signal (units of molecules per receptor) in response to two different ligands, ⟨*n /± σ* ⟩ _*n*_, as a function of the number of proofreading steps. Here we have set *k*_on_*c/k*_*p*_ = 1, *k*_*f*_ */k*_*p*_ = 1, *k*_*p*_*t* = 100, *k*_off,1_*/k*_*p*_ = 3 and *k*_off,2_*/k*_*p*_ = 1.5. Our choice of parameters emphasises the point that noise in the signals grows faster (with *N*) than the mean signals separate. C) The specificity as a function of of proofreading strength *g* and number of proofreading steps *N*, using the same parameters as (A).

The SNR also reveals that the output distributions from different ligands increasingly overlap as the number of proofreading steps increases, despite the increased separation of their means, as shown in Fig. 2B. The difference in the mean outputs in response to each ligand (blue and orange lines) increases with *N*, illustrating the classical proofreading effect. However, despite this increasing separation between the means, the output distributions become progressively more overlapped [colored envelopes represent 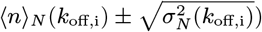. The SNR therefore already provides an indication that the noise associated with proofreading makes ligand resolution harder. In the next section we formulate receptor specificity in a precise mathematical manner and show that the intrinsic noise of the KPR scheme poses a challenge for KPR as a mechanism for enhanced specificity.

### C. Proofreading decreases specificity

To give a precise meaning to the problem of ligand resolution, and thereby quantify the effects of noise on ligand resolution, we frame it as a problem of determining the ligand identity based on the observed receptor output, *n*. This entails statistical estimation of *k*_off_ using an estimator function 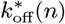. For Normally distributed outputs considered here, choosing 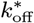 to be the maximum likelihood estimator (MLE) — that is, the estimate for which *P* (*n*|*k*_off_) is maximized — guarantees an unbiased estimate (i.e.,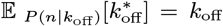) with a minimum variance over different estimators (35, 50, 60). At long times, the MLE is well approximated by the inverse function ⟨*n*⟨ ^−1^(*k*_off_) obtained by solving Eq. 1 for *k*_off_ (53, 60).

As illustrated in Fig. 1C, the distribution of receptor outputs, *P* (*n*|*k*_off_), gives rise to the distribution of the estimates of *k*_off_, 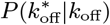 because the (deterministic) estimate 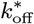 depends on *n* sampled from the distribution *P* (*n*|*k*_off_) (53, 60). Since the estimator is unbiased, 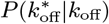 is centered at the correct estimate, 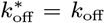. The width of the distribution of estimates depends on both the width of the output distribution, *P* (*n*|*k*_off_), and the estimator function, 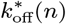, as illustrated in Fig. 1C. The Cramér-Rao lower bound provides an expression for the minimum expected variance of 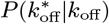 as: 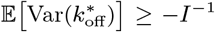 where 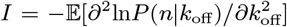 is the Fisher Information and all expectations are taken over *P* (*n*|*k*_off_) (60). Intuitively, the more information about *k*_off_ that is contained in *P* (*n*|*k*_off_), the lower is the variance in the estimate 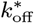. For a Normally distributed receptor output, *n*, the Fisher Information can be calculated as (53, 60):

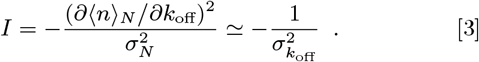

For sharply peaked output distributions the distribution of estimates, 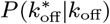, is well approximated by 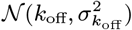, where 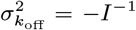 is the estimate variance. The estimate distribution for general *N* has variance:

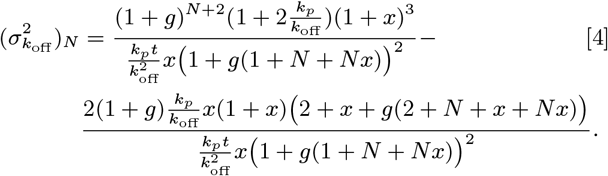

For strong proofreading the variance of the estimate scales as 𝒪 (*g*^*N*^). We and others have recently quantified receptor specificity by the ratio 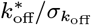, which is similar to the SNR but measures the relative width of the *estimation* distribution, 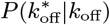 (48, 53). The specificity ratio is inspired by the measure of accuracy of ligand concentration sensing first calculated by Berg and Purcell (34). Specificity is a critical prerequisite for good ligand resolution because greater specificity allows the ligand affinities to be closer together without their estimates being statistically indistinguishable.

We demonstrate in Fig. 2C that, except for low *N* in the weak proofreading regime, *g*; ≲ 1, the specificity (calculated from Eq. 4) decreases with increasing number of proofreading steps, *N*. Notably, this detrimental effect of proofreading persists throughout the regime where weakly binding ligands are strongly proofread but strongly binding ligands are not (corresponding to 1 < *g* < 10 for the weakly binding ligand in Fig. 2C), which is the regime where KPR is commonly invoked - such as in TCR signaling. The decrease in specificity at large *N* is a consequence of the increased relative noise in the receptor output as proofreading steps are added (see Fig. 2B). Note that we interpolate the results for non-integer *N*, which can be thought of as the effective number of proofreading steps for imperfect proofreading such as when the proofreading steps are not perfectly irreversible (14). Specificity is maximized for low but nonzero values of *N* in the weak proofreading regime because of a balance between the detrimental effect of molecular noise and the beneficial effect of increased sensitivity to changes in *k*_off_ arising from the term |∂ ⟨*n)*_*N*_⟩ */*∂*k*_off_| in Eq. 3. At very small *g* the receptor spends effectively no time in the proofreading states and the specificity is decreased again. The balance of these effects results in the maximization of the specificity for low but nonzero values of *N* in the weak proofreading regime; the location of this maximum in (*g, N*)-space also depends on the ligand concentration (see Fig. S1 C). However, the enhancement of specificity compared to a non-proofread receptor is very moderate (less than 10-fold for any *N* with *ζ* = 0.1, Fig. S1 G) and is substantially lower than the classical and commonly invoked exponentially strong enhancement in the strong proofreading regime. Thus, the fact that specificity is only enhanced in this regime casts a new light on the performance and the purpose of KPR in such applications.

Another approach to quantify the effects of noise on ligand discrimination is through the mutual information, ℐ (*n, k*_off_), between the input variable (ligand affinity), *k*_off_, and the receptor output, *n* (18, 22, 59). Although not the main thrust of this paper, it has been shown that mutual information is closely related to the Fisher Information which we considered in this section (66, 67). As shown in the Supporting Information (S.I.) Section A, proofreading also decreases the discrimination when quantified through the mutual information.

We have shown that both strong proofreading and a large number of proofreading steps generally lead to very noisy estimates for a single ligand affinity. The greater the spread in the estimate distribution for a single ligand (i.e., the specificity), the less well resolved two different ligands are expected to be. In the next section we give this notion a precise mathematical meaning by constructing a resolution metric for two ligands.

### D. A metric for two-ligand resolution

While the SNR and the specificity are useful for quantifying noise in KPR performance both metrics describe the signaling statistics for only a single ligand. We now extend the idea of specificity to the problem of discriminating between two ligands that are are distinguished by their respective ligand-receptor dissociation rates *k*_off,1_ and *k*_off,2_. The differences in the affinity between the ligands is characterized by the ratio *ζ* = *k*_off,2_*/k*_off,1_ (we define *ζ*∈ (0, 1] so that *k*_off,1_ *> k*_off,2_). An appropriate resolution metric should encapsulate how well the difference in affinity between the two ligands, *ζ*, can be resolved in the presence of noise in the output distribution of each ligand.

The standard metric for ligand resolution in the KPR model, first used by Hopfield and Ninio, uses the ratio of the mean outputs for the two ligands (1–3):

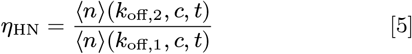

where ⟨*n* (*k*_off,*i*_⟩, *c, t*) is the mean signal produced by time *t* by the receptor in response to a ligand with dissociation rate *k*_off,*i*_ at fixed concentration *c*. In our convention *ζ* < 1 and *η*_HN_ ≥1 since the weaker binding ligand forms less long-lived ligand-receptor complexes and therefore produces less output. The larger *η*_HN_, the better these ligands can be discriminated. For concreteness, we choose *ζ* = 0.1 here and throughout this paper as informed by the typical value for self-peptides versus weak antigens to be separated by the TCR (3, 12, 14, 18, 68). As discussed above, the metric *η*_HN_ relies only on the average receptor outputs and does not account for the noise in the output distributions which we have shown to be so important for ligand discrimination.

To take the noise into account, a resolution metric must quantify the distance between the two estimate distributions 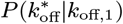 and 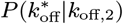. Intuitively, two distributions are well separated when their means are separated by more than the sum of their standard deviations (e.g., see Fig. 1B). This intuition is encapsulated in the Fisher’s linear discriminant (69, 70), based on which we define our ligand resolution metric:

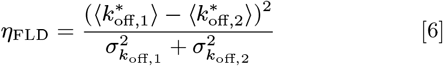

where 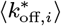 is the mean estimate for *k*_off_ generated by ligand with *k*_off,*i*_, and 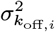 is the variance about this mean. The averages in Eq. 6 are taken over *P* (*n*|*k*_off,*i*_), and we assume an unbiased estimator so that 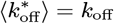 (60). Note that in the limit of similar ligands we have *k*_off,2_ ≈ *k*_off,1_ + *δk*_off_ and 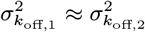, so that:

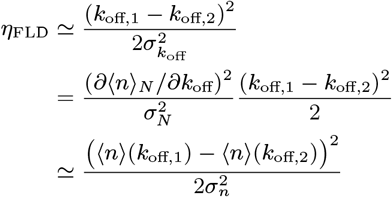

where we have used the Fisher Information of (Eq. 3) and in the last line we have interpreted d*k*_off_ = *k*_off,1_ −*k*_off,2_. Thus, the Fisher linear discriminant formulated in *k*_off_-space is very closely related to the equivalent formulation in *n*-space that conveys the intuitive meaning about the ability to separate the two output distributions. For both *η*_HN_ and *η*_FLD_, a higher value indicates higher resolution. However, we show in the following section that KPR has severely restricted benefits for ligand discrimination, relative to a non-proofreading receptor, when noise is accounted for in quantifying the resolution.

### E. Increasing the number of proofreading steps does not enhance discriminatory power

For a KPR receptor with *N* proofreading steps, the Hopfield-Ninio resolution (Eq. 5) is:

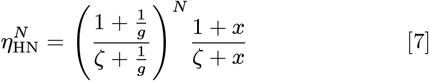

using *x* = *k*_on_*c/k*_off,1_, *g* = *k*_off,1_*/k*_*f*_ is the proofreading strength for the weaker binding ligand, and *ζ* = *k*_off,2_*/k*_off,1_. In the limit of strong proofreading (1*/g* → 0) and low concentration (*x* ≪ 1), the resolution described by Eq. (7) scales as 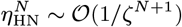. At high concentrations (*x ≫* 1) and strong proofreading the resolution scales as 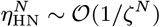 due to the saturation of the receptor by the ligand (Section 2C). Thus, the enhancement of the resolution by *N* proofreading steps is given by:

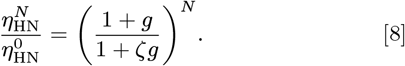

In the strong proofreading regime this simplifies to 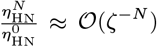, while in the weak proofreading regime this simplifies to 𝒪 (1). The behavior of *η*_HN_ expresses the well known result that in the strong proofreading regime, additional proofreading steps exponentially enhance mean resolution relative to a non-proofreading receptor. However, when noise is included in the resolution metric, as in *η*_FLD_, increasing the number of steps does not necessarily increase the discrimination strength because the separation of the means ⟨*n)*(*k*_off,i_) grows slower with *N* compared to the variances 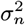, obscuring the distinctions between ligands.

A closed form expression for the resolution of the KPR receptor using the metric 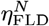 is cumbersome (provided for completeness in S.I.), but can easily be evaluated numerically. In Fig. 3A we show how *η*_FLD_ depends on the number of proof-reading steps, *N*, and proofreading strength, *g* (fixing *k*_*f*_ = 1 so that *g* = *k*_off_, *k*_on_*c* = 0.1, and *ζ* = 0.1). Unlike *η*_HN_, here an intermediate proofreading strength for the weaker binding ligand [*g* ∼ *𝒪* (1)] and a few proofreading steps yield the greatest resolution, similar to the the specificity in Fig. 2C, reflecting the mathematical similarity of the two metrics (discussed in Section D).

Although the location of the maximum of the resolution, *η*_FLD_, depends on the ligand concentration *c* (see Fig. S1 D-E), the maximum is attained at weak proofreading strengths (*g* < 1 for both ligands) for all concentrations (see inset of Fig. S1 D). The commonly considered context for KPR is where weakly binding non-agonists are suppressed by proofreading (1 < *g*_weak_ < 10) while strongly binding agonists are hardly suppressed (0.1 < *g*_strong_ < 1) (12, 14). In this biologically important regime, denoted with a dashed line in Fig. 3A, the resolution monotonically decreases as proofreading steps are added.

We visualise the resolution enhancement relative to the non-proofreading receptor by the ratio 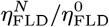 in Fig. 3B, which shows that the resolution does not improve with additional proofreading steps. Instead, the ratio scales inversely with *N* as *N* becomes large (green lines in Fig. 3B). While there can be moderate resolution enhancement for a few proofreading steps in the regime of weak proofreading, this enhancement is always ablated in the large *N* limit since the ratio scales as

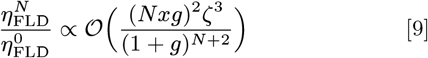

**Fig. 3.**
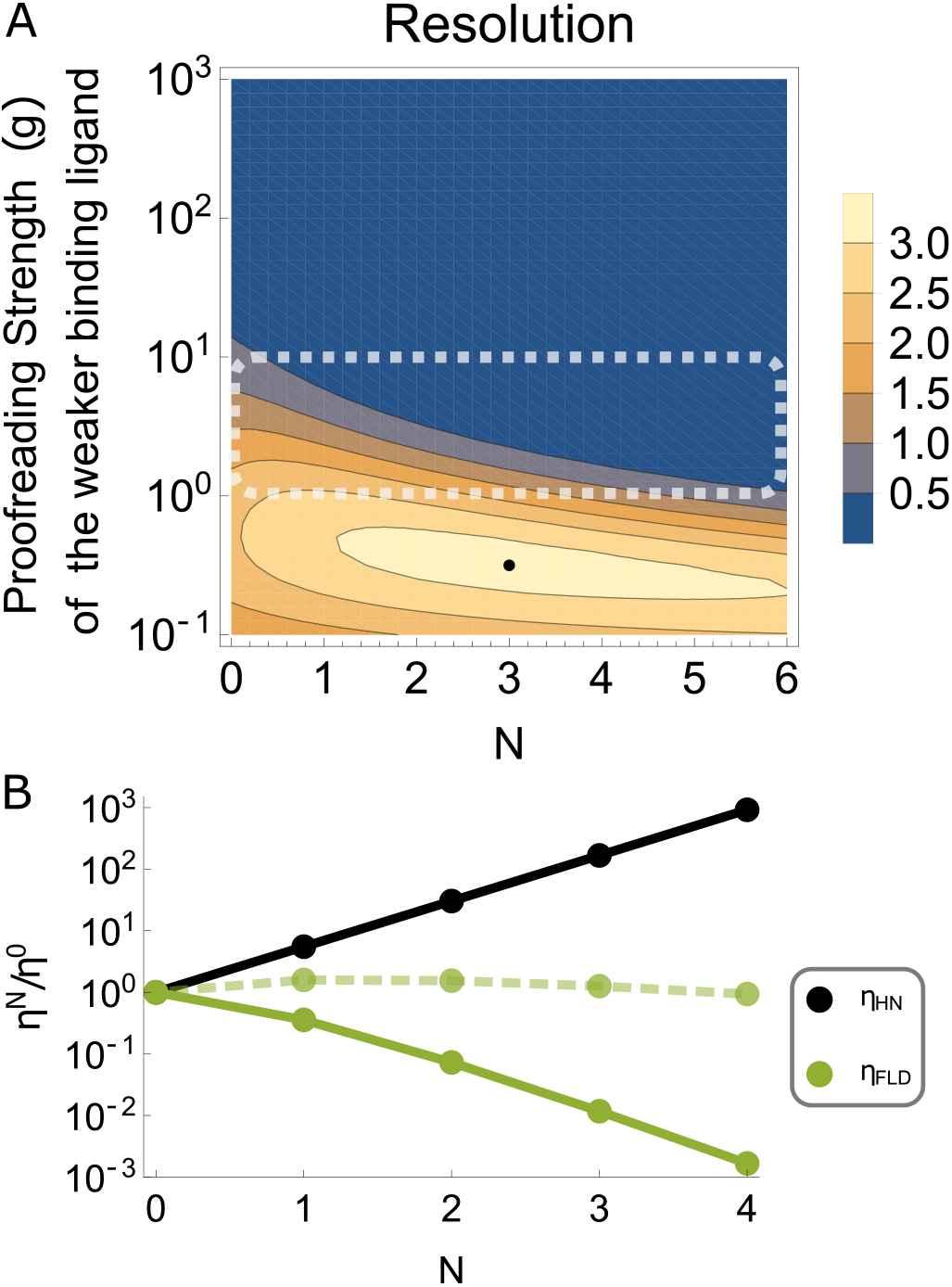
Kinetic proofreading does not enhance resolution. A) The resolution of a KPR receptor, as measured by *η*_FLD_ (Eq. 6), with *k*_on_ *c/k*_*p*_ = 0.1, *k*_*p*_*t* = 100, *k*_*f*_ */k*_*p*_ = 1, and *ζ* = 0.1. The resolution has a single maximum, indicated by the black point. The dashed lines indicate the biologically relevant scenario where weakly binding non-agonists are suppressed by proofreading but strongly binding agonists are not. B) The fold-change in *η*_FLD_ and *η*_HN_ achieved by KPR as a function of the number of proofreading steps, *N*, at *g* = 10 (solid green line), i.e., *k*_*f*_ */k*_*p*_ = 1, *k*_off,1_*/k*_*p*_ = 10 & *k*_off,2_*/k*_*p*_ = 1) or *g* = 1 (dashed green line), *k*_off,1_*/k*_*p*_ = 1 & *k*_off,2_*/k*_*p*_ = 0.1 and *k*_on_*c/k*_*p*_ = 1. Other parameters are the same as (A).

which tends to zero for *N* → ∞ — opposite to the scaling of 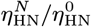 shown in Fig. 3B (black line). The resolution enhancement for a broad range of proofreading strengths is provided in the S.I. (Figure S1 G), which also shows that the fold-change in resolution given by 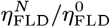 is significantly less than that indicated by 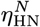.

The resolution at large *N* decreases because the relative noise of the receptor output is always increased by increasing the number of proofreading steps (see Fig. 2A). In principle, the molecular noise of the receptor could be reduced in different variants of the KPR mechanism. However, in the next section we demonstrate that even in models with minimal noise, the inherent stochasticity of the KPR mechanism still results in diminished resolution capacity compared to a non-proofread receptor (see also S.I. Section E).

### F. Alternative metrics and models of noise reveal intrinsic disadvantages of proofreading

#### F.1. Low noise models

The KPR model considered so far has several sources of noise. These include the intrinsic noise inherent to the probabilistic nature of the sequential state transitions during proofreading, the variability of the residence times in different states, and the noise in the production of the receptor output *n*. Furthermore, for mathematical tractability all rate processes have been assumed to be Poisson processes with exponentially distributed reaction times.

In this section, we investigate alternative KPR schemes with diminished noise to show that our results are not an artefact of a particular choice of the (high noise) model. In the first scheme (see Section D of the S.I.), to understand the role of the noise in the production of the output variable, we consider a KPR receptor which produces bursts of *β* output molecules immediately upon arriving in the active state (*N* + 1^*th*^) state. This model can serve as a representation of a fast signaling output burst by a G-protein coupled receptor (53, 71, 72) Unlike a Poisson rate process with rate *k*_*p*_, this burst process does not add extra noise associated with the output generation. The mean number of the output molecules, ⟨*m*⟩, produced by this burst model is

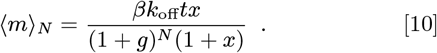

The variance in the number of output molecules as a function of *N* is derived in the Section D of the S.I..

In the second alternative scheme, we investigate the role of variance in the residence time in the active state of the receptor. To this end, in addition to the burst output the residence time in the active state, rather than obey an exponential distribution, is a constant *τ* = 1*/k*_off_ for each ligand. Furthermore, in the S.I. we present another version of this model where all transition times between states are *δ*-rather than exponentially distributed. This hybrid model has been investigated using stochastic simulations (see Methods).

The results of these minimal noise models are shown in Fig. 4A. The burst model shows a significant decrease in the resolution for the addition of a single proofreading step and a steady decline as *N* increases. The model with a fixed residence time is less noisy and consequently shows better resolution than either burst or Poisson types of output with exponentially distributed times. Yet, both minimal noise models exhibit decreasing discriminating power with increasing number of proofreading steps, similar to the original KPR model(Eq. 9). As above, the resolution of these models is worse in the strong proofreading regime (*g* = 10, solid line) compared to the weak proofreading regime (*g* = 3, dashed line).

#### F.2. Alternative metrics of ligand resolution

In the previous sections we have assumed that the measurement time is sufficiently long so that the distribution of the output, *n*, is well approximated by a normal distribution. However, in some situation – notably the problem of antigen detection by TCR – it is possible that a cell must determine a ligand identity very quickly, ruling out the possibility for many binding-unbinding events to occur during the measurement time. We investigated this regime using stochastic simulation (see S.I. Section G), and the results are shown in Fig. 4B, which shows that even at very short times *η*_FLD_ decreases with *N*, as seen for long measurement times.

Furthermore, in some cases – including the problem of TCR discrimination – it can be argued that the primary purpose of KPR for a T cell is the maximal suppression of false positive errors from the weakly binding ligand (even at the expense of the somewhat increased false negative rate for the strongly binding ligand). Spurious activation of the immune system by weak binders - which carries the potential of substantial damage to the host - may be more dangerous to the host compared to false negative errors of occasional non-detection of the strongly binding ligand because the latter can be compensated for by other clonal cells that identify the non-self antigen (73, 74).

We quantify these notions through a metric defined as the ratio of the probability of false activation to the probability of all activation events. We assume that activation occurs when the output variable *n* crosses an activation threshold *n*\*, with probability *P* (*n*(*k*_off, weak_, *c*) *> n*\*). The threshold is set to a value such that the probability of activation by the strongly binding ligand is 10%: *P* (*n*(*k*_off, strong_, *c*) > *n*\*) = 0.1 (the results are robust with respect to the specific choice of this value). In Fig. 4C we show the probability ratio of false activation to the total activation as a function of the number of proofreading steps This plot reveals that the ratio is minimized for a small number of proofreading steps and increases as *N* is increased further. The location of this minimum shifts to larger *N* as the proofreading strength decreases, and the minimum shifts to smaller *N* as the proofreading strength increases. We show the ratio for *g*_weak_ = 10 and *g*_weak_ = 3, which represent a typical range of proofreading strengths for the TCR according to recent experimental measurements (12–14).

Overall, our results demonstrate that KPR has severely limited benefits for ligand discrimination, only providing moderate enhancement in ligand resolution in the non-classical regime of fast proofreading and a few proofreading steps.

## 3. Discussion

In this paper we have shown that intrinsic molecular noise greatly affects the KPR-based ability of a receptor to discriminate between ligands, and that resolution metrics must account for this noise to avoid misleading interpretations of the effects of KPR. To capture the role of the noise, we used the metric *η*_FLD_ (Eq. 6) which, unlike the classical metric *η*_HN_ (Eq. 5), incorporates information not only about the means but also about the variances of the distributions of the estimated ligand identity, 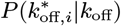. Taking the noise into account reveals that the benefits of proofreading for ligand discrimination are severely restricted to a moderate effect at low number of proofreading steps and low proofreading strength.

The central result of the classical analysis of KPR (based on *η*_HN_) posits that the discrimination between two ligands improves with the number of proofreading steps as 𝒪 (*ζ*^−(*N*+1)^) in the strong proofreading regime, where *ζ* = *k*_off,2_*/k*_off,1_ is the ratio of the affinities of the two ligands to be distinguished. By contrast, in Sections 2 B-E we demonstrated that proofreading amplifies the effect of intrinsic molecular noise in both the signaling state occupancy and the downstream output, thereby making ligand discrimination more difficult. In all regimes, for large *N*, the resolution enhancement, 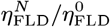, scales inversely with *N* and tends to zero as *N* → ∞.

One of the most common biological signaling systems where the KPR model has been invoked is T cell discrimination between foreign antigens and self peptides (43). These applications of the KPR model commonly focus on the scenario where the number of proofreading steps is large (18, 21, 27, 61) and the weakly binding self peptides are strongly proofread and attenuated, because this is where the proofreading is most effective according to the classical, noiseless, HN metric. However, the large *g*, large *N* regime is also the regime where we showed the receptor output distribution to be most noisy leading to the degradation in discrimination performance (Section 2B. While experimental measurements of KPR parameters in biological systems have remained difficult, recent work has started to provide experimental estimates of the effective number of proofreading steps for the TCR, estimated to be *N* ≈3 (13, 14), and estimates of the proofreading rate have been found to be *k*_*f*_ ≈ 0.3*s*^−1^ (14). Thus, for a typical agonist binding time of 1 - 10 seconds, a typical non-agonist binding time of 0.3 - 3 seconds, and taking *k*_off_ = *τ* ^−1^ for the binding time *τ* (12), the TCR operates with moderate proofreading strength: 0.3 < *g* < 3 for the strongly binding ligand and 1 < *g* < 10 for the weakly binding ligand. Weakly binding ligands may be proofread even more strongly according to recent re-measurement of ligand affinities for the TCR (14). These experimental measurements of *N* and *g* for the TCR are thus consistent with the conclusions of our theoretical framework, although full comparison of the theoretical predictions with the detailed the experimental data is outside the scope of the present work and will be investigated in the future.

**Fig. 4.**
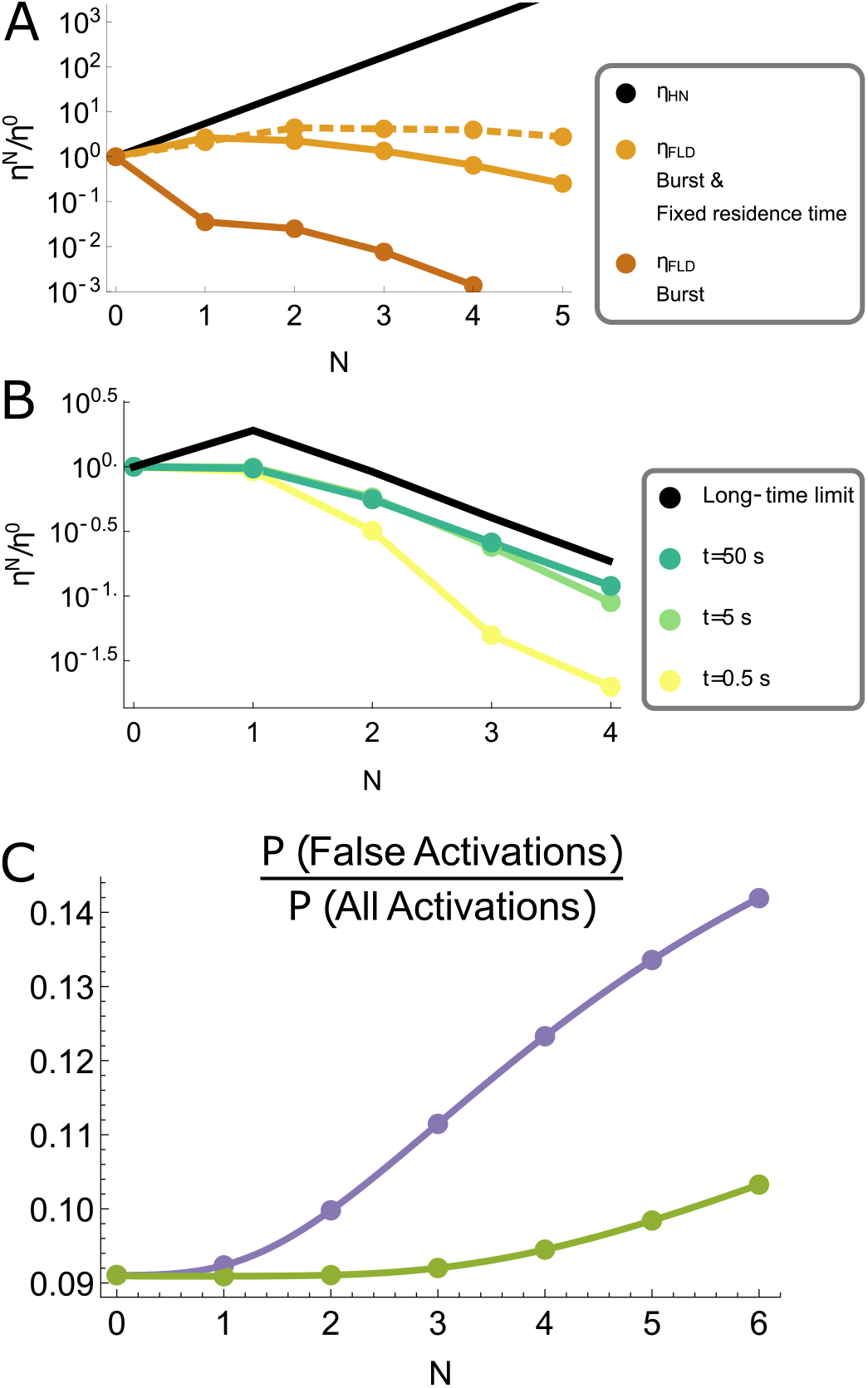
Classification performance is best for a small number of weakly proof-read steps. A) The fold-change in resolution achieved by KPR as a function of the number of proofreading steps, *N*, but for the alternative KPR schemes discussed in Section 2 F: the burst type output model (red) and the burst model with fixed residence time, *τ* = 1*/k*_off_, in the final receptor state (yellow). For the fixed residence time model, resolution is shown at proofreading strengths *g*_weak_ = 3 (dashed) and *g*_weak_ = 10 (solid). The burst size is 10, *k*_off,1_*/k*_off,2_ = 0.1, *k*_*f*_ */k*_on_*c* = 2(6.7) for the strong (weak) proofreading, and *k*_on_ *ct* = 500. B) The ligand resolution measured as 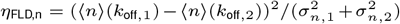 of the fully stochastic KPR receptor model at short times. The Hopfield-Ninio resolution is provided for reference (black). The resolution increases at longer times because receptor output accumulates and thereby reduces shot noise that otherwise obscures ligand resolution. However, the resolution enhancement decreases for *N >* 1 (more than one proofreading step), the same trend as was observed at long times (see Fig. 3C). C) The probability of erroneous activation relative to the total probability of activation, as a function of the number of proofreading steps *N*, for proofreading strength *g* = 10 (purple) versus *g* = 3 (green). Parameters: *k*_on_*c/k*_*p*_ = 1, *k*_*p*_*t* = 100, *k*_off,1_*/k*_*p*_ = 1 & *k*_off,2/*k_p_*_ = 0.1, and *k*_*f*_/*k*_*p*_ = 0.1(0.3) for the stronger (weaker) proofreading strength. Interpolation between integer *N* made using Eqs. 1&2.

The limited utility of KPR in ligand discrimination and identifiability stems from its two inherent properties: attenuation of the occupancy of the signaling state, and the accompanying amplification of the effects of noise with each additional proofreading step. The problems posed by signal attenuation were already noted by McKeithan (3), who proposed a variant KPR scheme in which the receptor is stabilized in the final signaling state. Such stabilization can be introduced into our model by reducing the unbinding rate from the final proofreading state by a factor *α* < 1 (see Methods: Eq. 11). The mean signal for a stabilized KPR receptor is

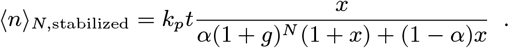

While stabilizing the signaling state does increase the mean signal, the standard deviation still scales like 𝒪 (*g*^−*N/*2^) so that the increased noise in the signal distribution still outpaces the increased difference in signals as proofreading steps are added (see also Fig. S2C).

The results of this paper are not an artefact of a particular choice of the noise model. The variability of the signaling output arises from several different components. The first two are the Poisson noise of the output production in the signaling state and the variability of the residence and arrival times to the signaling state. The third is the probabilistic nature of the binding-unbinding events and state transition steps, which are intrinsic to the proofreading mechanism. As shown in Section 2F, the results of this paper - the decrease of the resolution capacity with stronger proofreading and increasing number of proofreading steps - hold true even in the minimal noise models where the signaling output is produced in discrete bursts and the receptor residence times are not exponentially but rather *δ*-distributed. Our results also hold true for fast decision-making processes which occur at short times when multiple binding-unbinding events are unlikely.

Our results rest on the appropriate choice of resolution metric *η*_FLD_ that is widely used in statistical inference as a measure of the estimate quality and discriminability. However, in different biological situations, a different metric can be more appropriate - such as the fraction of the false positive activations. As shown in Section 2F, the results of this paper are not an artefact of a particular metric choice, and hold also for different measures of the discrimination quality; the fraction of false positives generally decreases with an increase in the number of proofreading steps, except for a moderate minimum at low *N* at low proofreading strength.

It is important to distinguish these results from other works that have sought to balance ligand discrimination performance with the extrinsically imposed costs of proofreading such as input energy and decision speed, finding that different choices of the extrinsic costs optimize the proofreading at at different numbers of proofreading steps (22, 39, 44–46, 75). These approaches typically posit that larger *N* will enhance ligand discrimination so long as the extrinsic costs can be met. By contrast our result indicate that the discrimination capacity itself decreases with the proofreading strength due to the intrinsically stochastic nature of KPR.

Our results naturally lead to a question: if KPR commonly does not enhance the resolution of a sensory system, what other purpose might it serve in receptor systems like the TCR? One possibility is that KPR might enable multiplexing of signals in receptor mixtures by producing multiple statistically independent outputs from different proofreading states (53, 76, 77). These outputs can be used to infer the presence of multiple ligands in a mixture (53), convey information about ligand concentration or other environmental conditions in addition to ligand identity (77), or perform other complex signaling processing tasks (78). Mounting experimental evidence indicates that the TCR (as a case study for KPR) may indeed produce multiple intracellular second messengers which may enable T cells to detect antigen quantity alongside antigen quality (78–82). Our results lay the ground for further theoretical work aimed at the understanding the effects of noise in multi-output receptor signaling pathways (78, 83–85).

## Materials and Methods

The KPR receptor states are enumerated by index *i*, with *i* = 0 representing the unbound state and *i* = *N* the signaling state. The system is fully characterized by the pair of state variables (*i, n*) representing the current receptor state and number of copies of the downstream signal molecule respectively. The probability of the system to be in the state *i* at time *t* is 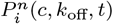 which depends on the ligand concentration *c* and the unbinding rate *k*_off_. The dynamics of this probability distribution is described by the Master equation

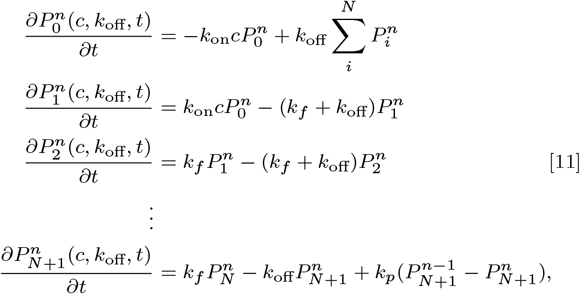

where *k*_on_ association rate of the free ligands with an unoccupied receptor, *k*_*f*_ is the rate at which the receptor transitions from state *i* to state *i* + 1, *k*_*p*_ is the rate of signal production from the terminal receptor state; we assume that the ligands are in a large extracellular volume so that *c* remains constant. A stabilized KPR receptor mentioned in the Discussion has *k*_off_ reduced by a factor *α* in the last line of Eq. 11. For the burst model of Section 2F the terms involving *k*_*p*_ are not present and the term 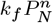 becomes 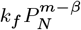 (see also S.I.). The mean and variance for the burst model are s^*N*^olved in a similar manner to that of the phosphorylation model, described below.

The probability distribution of the output signal is 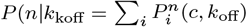. Accordingly, the mean downstream output signal *n* is

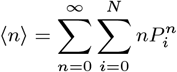

Taking the derivative with respect to time and using the master equation above, the summation over *P*_*i*_ reduces this expression to

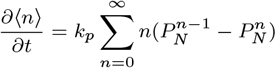

Re-indexing the term 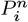 leads to

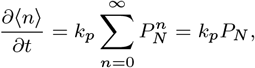

where *P*_*N*_ is the steady-state occupancy probability for the *N* ^th^ receptor state. The steady-state receptor occupancy probabilities *P*_*i*_(*c, k*_off_) can be obtained by algebraically solving the system of equations resulting from setting all time derivatives and *k*_*p*_ to zero in Eq. 11 and summing over the state variable *n* in 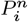. Finally, the mean signal downstream of an *N*-state KPR receptor is

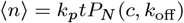

Using the expression for *P*_*N*_ for a general *N*-step proofreading receptor that can be obtained from the Master equation Eq. 11 yields Eq. 1 for the mean signal (3): The case of *N* = 0 is a non-proofreading receptor. The same procedure can be used to compute the average output from the G-protein type KPR receptor, with the main difference being that the Master equation yields the relation ∂ ⟨*m*⟨ /∂*t* = *k*_*f*_ *P*_*N*−1_ for *N* > 0 and ∂ *m* / ∂*t* = *k*_on_*cP*_0_ for *N* = 0.

The variance in the downstream signal, 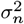, can be obtained from the second moment of the signal using the identity 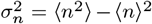. The second moment is defined as

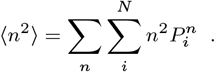

Once again, taking the derivative with respect to time and substituting the Master equation yields a closed form expression:

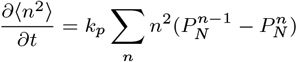

Re-indexing the term 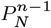 leads to:

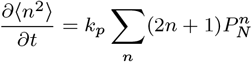

A closed form expression for the term 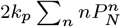 can be obtained by solving the associated set of inhomogenous first order differential equations:

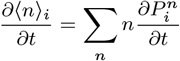

for *i* ∈ [0, *N*]. Substituting the resulting expression for *n)*_*i*_ into the differential equation for 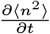 and integrating over time yields an expression for the second^∂^m^*t*^ oment, which can then be used to find the variance 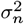. Using closed form expressions for *P*_*N*_, the variance of the downstream signal is, in the long time limit min(*k*_off_*t, k*_*f*_ *t*) → ∞, Eq. 2. For the G-protein type KPR receptor the Master equation yields 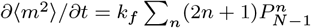, but the procedure is otherwise the same.

We view the T cell sensing problem as fundamentally a classification task based on estimating the ligand dissociation rate *k*_off_. A maximum likelihood estimator (MLE) for *k*_off_ can be constructed from the distribution 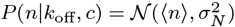. At long times the limit of large numbers applies, making the distribution *P* (*n*| *k*_off_ narrowly peaked about its mean so that this MLE is well approximated by (35,36.

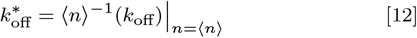

The inverse function ⟨*n* ⟩^−1^ is easily obtained by simple algebraic manipulation of Eq. 1 for all choices of *N* considered in this paper.

Variation in the observed signal *n* results in variation of the estimate for *k*_off_. We define a particular trajectory of the random variable *n*, which depends on the ligand dissociation rate, as *n*(*k*_off_, *t*). An expression for the estimation variance can be obtained by taking a first-order Taylor expansion of *n* about *k*_off_:

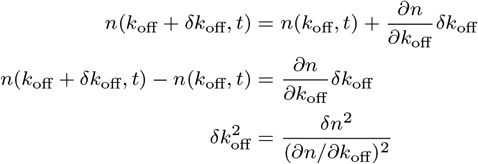

where in the last line we have defined *δn* = *n*(*k*_off_+*δk*_off_, *t*)−*n*(*k*_off_, *t*) and squared the expression. Averaging this expression over the (time-dependent) signal distribution 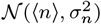and interpreting quantities like ⟨*δ·*^*2*^⟩ as variances yields a simple form for the variance of the estimate distribution:

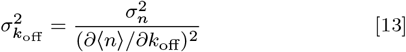

This expression can be more rigorously derived by using the Cramér-Rao lower bound, which states that the variance in an unbiased estimator 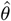 is bounded below by

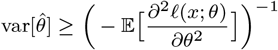

where *ℓ*(*x*; *θ*) is the log-likelihood of observation *x* given parameter *θ* (60). In the long-time limit of the KPR model the likelihood for *n* is well-approximated by a Gaussian, which allows one to write the right hand side of the Cramér-Rao lower bound as the inverse Fisher Information (53, 60).

Means and variances for the semi-deterministic KPR models and the stabilized KPR receptor model were computed by Gillespie simulation (86). The fixed residency time in the final proofreading state was simulated by replacing the random time step in the Gillespie algorithm with a deterministic time step *τ* = 1*/k*_off_. For the minimal noise model treated in the S.I., the receptor either transitions to the next proofreading state or the ligand unbinds the receptor. The probability of either event is *k*_*f*_ */*(*k*_*f*_ + *k*_off_) and *k*_off_*/*(*k*_*f*_ + *k*_off_) respectively. Once the next reaction is selected in the simulation, the time for that reaction to happen is determined as either 1*/k*_*f*_ or 1*/k*_off_, respectively.

## ACKNOWLEDGMENTS

The authors are indebted to Golan Bel, Sid Goyal, Matthew Smart, Jeremy Rothschild, and numerous colleagues in the field for illuminating discussions. The authors acknowledge the support of the Natural Sciences and Engineering Research Council of Canada (NSERC) through a Discovery Grant RGPIN 402591 to A.Z. and a PGS-D Graduate Fellowship to D.K.

## References

1. J Hopfield, Kinetic Proofreading: A New Mechanism for Reducing Errors in Biosynthetic Processes Requiring High Specificity. PNAS 71, 4135–4139 (1974).

2. J Ninio, Kinetic amplification of enzyme discrimination. Biochimie 57, 587–595 (1975).

3. TW Mckeithan, Kinetic proofreading in T-cell receptor signal transduction. Immunology 92, 5042–5046 (1995).

4. KA Hogquist, SC Jameson, MJ Bevan, Strong agonist ligands for the T cell receptor do not mediate positive selection of functional CD8+ T cells. Immunity 3, 79–86 (1995).

5. SM Alam, et al., Qualitative and quantitative differences in T cell receptor binding of agonist and antagonist ligands. Immunity 10, 227–237 (1999).

6. JT Reardon, A Sancar, Thermodynamic cooperativity and kinetic proofreading in DNA damage recognition and repair. Cell Cycle 3, 139–142 (2004).

7. G Altan-Bonnet, RN Germain, Modeling T cell antigen discrimination based on feedback control of digital ERK responses. PLoS Biol. 3, 1925–1938 (2005).

8. C Torigoe, JR Faeder, JM Oliver, B Goldstein, Kinetic Proofreading of Ligand-Fc ϵ RI Interactions May Persist beyond LAT Phosphorylation. The J. Immunol. 178, 3530–3535 (2007).

9. PK Tsourkas, W Liu, SC Das, SK Pierce, S Raychaudhuri, Discrimination of membrane antigen affinity by B cells requires dominance of kinetic proofreading over serial engagement. Cell. Mol. Immunol. 9, 62–74 (2012).

10. MA Daniels, et al., Thymic selection threshold defined by compartmentalization of Ras/MAPK signalling. Nature 444, 724–729 (2006).

11. O Stepanek, et al., Coreceptor scanning by the T cell receptor provides a mechanism for T cell tolerance. Cell 159, 333–345 (2014).

12. G Altan-Bonnet, T Mora, AM Walczak, Quantitative Immunology for Physicists. Phys. Reports 849, 1–83 (2020).

13. DK Tischer, OD Weiner, Light-based tuning of ligand half-life supports kinetic proofreading model of T cell signaling. eLife 8, 1–25 (2019).

14. J Pettmann, et al., The discriminatory power of the T cell receptor. eLife 10, 1–42 (2021).

15. JA Owen, TR Gingrich, JM Horowitz, Universal Thermodynamic Bounds on Nonequilibrium Response with Biochemical Applications. Phys. Rev. X 10, 11066 (2020).

16. A Murugan, DA Huse, S Leibler, Speed, dissipation, and error in kinetic proofreading. Proc. Natl. Acad. Sci. 109, 12034–12039 (2012).

17. PS Swain, ED Siggia, The role of proofreading in signal transduction specificity. Biophys. J. 82, 2928–2933 (2002).

18. RS Ganti, et al., How the T cell signaling network processes information to discriminate between self and agonist ligands. Proc. Natl. Acad. Sci. United States Am. 117, 26020–26030 (2020).

19. H Mellenius, M Ehrenberg, Transcriptional accuracy modeling suggests two-step proofreading by rna polymerase. Nucleic Acids Res. 45, 11582–11593 (2017).

20. A Murugan, DA Huse, S Leibler, Discriminatory proofreading regimes in nonequilibrium systems. Phys. Rev. X 4, 1–8 (2014).

21. M Lever, PK Maini, PA Van Der Merwe, O Dushek, Phenotypic models of T cell activation. Nat. Rev. Immunol. 14, 619–629 (2014).

22. P François, G Altan-Bonnet, The Case for Absolute Ligand Discrimination: Modeling Information Processing and Decision by Immune T Cells. J. Stat. Phys. 162, 1130–1152 (2016).

23. WD Piñeros, T Tlusty, Kinetic proofreading and the limits of thermodynamic uncertainty. Phys. Rev. E 101, 1–8 (2020).

24. O Feinerman, RN Germain, G Altan-Bonnet, Quantitative challenges in understanding ligand discrimination by α β T cells. Mol. Immunol. 45, 619–631 (2008).

25. I Štefanová, et al., TCR ligand discrimination is enforced by competing ERK positive and SHP-I negative feedback pathways. Nat. Immunol. 4, 248–254 (2003).

26. OS Yousefi, et al., Optogenetic control shows that kinetic proofreading regulates the activity of the t cell receptor. eLife 8, 1–33 (2019).

27. WS Hlavacek, A Redondo, C Wofsy, B Goldstein, Kinetic proofreading in receptor-mediated transduction of cellular signals: Receptor aggregation, partially activated receptors, and cytosolic messengers. Bull. Math. Biol. 64, 887–911 (2002).

28. WS Hlavacek, JR Faeder, ML Blinov, AS Perelson, B Goldstein, The Complexity of Complexes in Signal Transduction. Biotechnol. Bioeng. 84, 783–794 (2003).

29. A Košmrlj, M Kardar, AK Chakraborty, Statistical physics of T-cell development and pathogen specificity. Annu. Rev. Condens. Matter Phys. 4, 339–360 (2013).

30. V Galstyan, K Husain, F Xiao, A Murugan, R Phillips, Proofreading through spatial gradients. eLife 9, e60415 (2020).

31. D Sagi, T Tlusty, J Stavans, High fidelity of RecA-catalyzed recombination: A watchdog of genetic diversity. Nucleic Acids Res. 34, 5021–5031 (2006).

32. PR ten Wolde, NB Becker, TE Ouldridge, A Mugler, Fundamental Limits to Cellular Sensing. J. Stat. Phys. 162, 1395–1424 (2016).

33. P François, A Zilman, Physical approaches to receptor sensing and ligand discrimination. Curr. Opin. Syst. Biol. in press, 1–11 (2019).

34. HC Berg, EM Purcell, Physics of chemoreception. Biophys. J. 20, 193–219 (1977).

35. RG Endres, NS Wingreen, Maximum likelihood and the single receptor. Phys. Rev. Lett. 103, 1–4 (2009).

36. T Mora, Physical Limit to Concentration Sensing Amid Spurious Ligands. Phys. Rev. Lett. 115, 1–9 (2015).

37. DC Wylie, J Das, AK Chakraborty, Sensitivity of T cells to antigen and antagonism emerges from differential regulation of the same molecular signaling module. Proc. Natl. Acad. Sci. United States Am. 104, 5533–5538 (2007).

38. G Bel, B Munsky, I Nemenman, The simplicity of completion time distributions for common complex biochemical processes. Phys. Biol. 7 (2010).

39. AH Lang, CK Fisher, T Mora, P Mehta, Thermodynamics of statistical inference by cells. Phys. Rev. Lett. 113, 1–5 (2014).

40. l Rayleigh, XV. On the theory of optical images, with special reference to the microscope. The London, Edinburgh, Dublin Philos. Mag. J. Sci. 42, 167–195 (1896).

41. H Sun, Basic optical engineering for engineers and scientists. (SPIE, Bellingham, Washington), (2018).

42. P François, G Voisinne, ED Siggia, G Altan-Bonnet, M Vergassola, Phenotypic model for early T-cell activation displaying sensitivity, specificity, and antagonism. Proc. Natl. Acad. Sci. United States Am. 110, E888–E897 (2013).

43. NC Trendel, O Dushek, Mathematical modelling of t cell activation. Math. Comput. Exp. T Cell Immunol. 1, 223–240 (2021).

44. G Tkačik, AM Walczak, W Bialek, Optimizing information flow in small genetic networks. Phys. Rev. E - Stat. Nonlinear, Soft Matter Phys. 80, 1–18 (2009).

45. JD Mallory, AB Kolomeisky, OA Igoshin, Trade-offs between error, speed, noise, and energy dissipation in biological processes with proofreading. J. Phys. Chem. B 123, 4718–4725 (2019).

46. W Cui, P Mehta, Identifying feasible operating regimes for early t-cell recognition: The speed, energy, accuracy trade-off in kinetic proofreading and adaptive sorting. PLoS ONE 13 (2018).

47. W Bialek, S Setayeshgar, Cooperativity, sensitivity, and noise in biochemical signaling. Phys. Rev. Lett. 100, 1–4 (2008).

48. ED Siggia, M Vergassola, Decisions on the fly in cellular sensory systems. Proc. Natl. Acad. Sci. United States Am. 110 (2013).

49. K Kaizu, et al., The berg-purcell limit revisited. Biophys. J. 106, 976–985 (2014).

50. M Carballo-Pacheco, et al., Receptor crosstalk improves concentration sensing of multiple ligands. Phys. Rev. E: Stat. Nonlinear, Soft Matter Phys. 99, 1–14 (2019).

51. T Mora, I Nemenman, Physical limit to concentration sensing in a changing environment. Phys. Rev. Lett. 123, 198101 (2019).

52. V Singh, I Nemenman, Universal Properties of Concentration Sensing in Large Ligand-Receptor Networks. Phys. Rev. Lett. 124, 28101 (2020).

53. D Kirby, J Rothschild, M Smart, A Zilman, Pleiotropy enables specific and accurate signaling in the presence of ligand cross talk. Phys. Rev. E 103, 42401 (2021).

54. C Waltermann, E Klipp, Information theory based approaches to cellular signaling. Biochim-ica et Biophys. Acta - Gen. Subj. 1810, 924–932 (2011).

55. G Tkačik, AM Walczak, W Bialek, Optimizing information flow in small genetic networks. III. A self-interacting gene. Phys. Rev. E - Stat. Nonlinear, Soft Matter Phys. 85 (2012).

56. T Jetka, K Nienałtowski, T Winarski, S Błoński, M Komorowski, Information-theoretic analysis of multivariate single-cell signaling responses. PLoS Comput. Biol. 15, 1–23 (2019).

57. P Binder, ND Schnellbacher, T Hofer, NB Becker, US Schwarz, Optimal ligand discrimination by asymmetric dimerization and turnover of interferon receptors. Proc. Natl. Acad. Sci. United States Am. 118 (2021).

58. M Komorowski, DS Tawfik, The Limited Information Capacity of Cross-Reactive Sensors Drives the Evolutionary Expansion of Signaling. Cell Syst. 8, 76–85.e6 (2019).

59. Y Tang, A Hoffmann, Quantifying information of intracellular signaling: Progress with machine learning. Reports on Prog. Phys. 85 (2022).

60. SM Kay, Fundamentals of Statistical Signal Processing: Estimation Theory. (Prentice-Hall, Inc., Upper Saddle River, NJ, USA), (1993).

61. DM Britain, JP Town, OD Weiner, Progressive enhancement of kinetic proofreading in t cell antigen discrimination from receptor activation to dag generation. eLife 11 (2022).

62. H Lodish, et al., Molecular Cell Biology. (W.H. Freedman), 4th edition, (2000).

63. CC Govern, PRT Wolde, Optimal resource allocation in cellular sensing systems. Proc. Natl. Acad. Sci. United States Am. 111, 17486–17491 (2014).

64. M Lever, et al., Architecture of a minimal signaling pathway explains the t-cell response to a 1 million-fold variation in antigen affinity and dose. Proc. Natl. Acad. Sci. United States Am. 113, E6630–E6638 (2016).

65. F Kok, et al., Disentangling molecular mechanisms regulating sensitization of interferon alpha signal transduction. Mol. Syst. Biol. 16, 1–32 (2020).

66. XX Wei, AA Stocker, Mutual information, fisher information, and efficient coding. Neural Comput. 28, 305–326 (2016).

67. N Brunel, JP Nadal, Mutual information, fisher information, and population coding. Neural Comput. 10, 1731–1757 (1998).

68. M Aleksic, et al., Different affinity windows for virus and cancer-specific T-cell receptors – implications for therapeutic strategies. Eur J Immunol. 42, 3174–3179 (2012).

69. T Hastie, R Tibshirani, J Friedman, The Elements of Statistical Learning: Data Mining, Inference, and Prediction. (Springer), 2 edition, p. 745 (2009).

70. RA Fisher, The Use of Multiple Measurements in Taxonomic Problems. Annals Eugen. 7, 179–188 (1936).

71. D Wootten, A Christopoulos, M Marti-Solano, MM Babu, PM Sexton, Mechanisms of signalling and biased agonism in G protein-coupled receptors. Nat. Rev. Mol. Cell Biol. 19, 638–653 (2018).

72. SR Neves, PT Ram, R Iyengar, G protein pathways. Science 296, 1636–1639 (2002).

73. TC Butler, M Kardar, AK Chakraborty, Quorum sensing allows t cells to discriminate between self and nonself. Proc. Natl. Acad. Sci. United States Am. 110, 11833–11838 (2013).

74. F Baumgart, M Schneider, GJ Schütz, How t cells do the “search for the needle in the haystack”. Front. Phys. 7 (2019).

75. K Banerjee, AB Kolomeisky, OA Igoshin, Elucidating interplay of speed and accuracy in biological error correction. Proc. Natl. Acad. Sci. 114, 5183–5188 (2017).

76. P Li, MB Elowitz, Communication codes in developmental signaling pathways. Dev. (Cambridge) 146, 1–12 (2019).

77. G Minas, et al., Multiplexing information flow through dynamic signalling systems. PLoS Comput. Biol. 16, e1008076 (2020).

78. SR Achar, et al., Universal antigen encoding of T cell activation from high-dimensional cytokine dynamics. Sci. (New York, N.Y.) 376, 880–884 (2022).

79. H Teimouri, AB Kolomeisky, Relaxation Times of Ligand-Receptor Complex Formation Control T Cell Activation. Biophys. J. 119, 182–189 (2020).

80. G Voisinne, et al., T Cells Integrate Local and Global Cues to Discriminate between Struc-turally Similar Antigens. Cell Reports 11, 1208–1219 (2015).

81. I Eizenberg-Magar, et al., Diverse continuum of CD4+ T-cell states is determined by hierarchical additive integration of cytokine signals. Proc. Natl. Acad. Sci. 114, E6447–E6456 (2017).

82. N Miskov-Zivanov, MS Turner, LP Kane, PA Morel, JR Faeder, The duration of T cell stimulation is a critical determinant of cell fate and plasticity. Sci. Signal. 6 (2013).

83. R Wolchinsky, et al., Antigen-Dependent Integration of Opposing Proximal TCR-Signaling Cascades Determines the Functional Fate of T Lymphocytes. The J. Immunol. 192, 2109–2119 (2014).

84. CY Ma, JC Marioni, GM Griffiths, AC Richard, Stimulation strength controls the rate of initiation but not the molecular organization of TCR-induced signalling. eLife 9, 1–57 (2020).

85. A Adelaja, et al., Six distinct NFκB signaling codons convey discrete information to distinguish stimuli and enable appropriate macrophage responses. Immunity 54, 916–930.e7 (2021).

86. DT Gillespie, Stochastic Simulation of Chemical Kinetics. Annu. Rev. Phys. Chem. 58, 35–55 (2007).

