## Supporting Information for "Proofreading Is Too Noisy For Effective Ligand Discrimination"

### 4. Supporting Information

**A. Mutual Information of the KPR Receptor.** The mutual information (M.I.) is typically interpreted as a measure of the fidelity of information transmission through a system, with a higher M.I. indicating that more information is successfully received. A higher M.I. is also often used to claim better ligand discrimination. The mutual information is given by $\mathcal{I}(n, k_{\text{off}}) = H(n) - H(n|k_{\text{off}})$ where $H(n|k_{\text{off}}) = -\mathbb{E}_{k_{\text{off}}}[\int P(n|k_{\text{off}})\log_2 P(n|k_{\text{off}})dn]$, $H(n) = -\int P(n)\log_2 P(n)dn$, the receptor output is Normally distributed as $P(n|k_{\text{off}}) = \mathcal{N}\Big(\langle n\rangle_N, \sqrt{\sigma_N^2}\Big)$, and $P(n) = \int P(n|k_{\text{off}})P(k_{\text{off}})dk_{\text{off}}$. We follow Ganti et al. [18] and assume $P(k_{\text{off}})$ is log-normal distributed, lognorm($\langle \log k_{\text{off}}\rangle = 0, 1$). Our results are robust to variation in this prior probability, $P(k_{\text{off}})$. The mutual information is plotted as a function of the number of proofreading steps, $N$, for different proofreading rates $k_f$ in Figure S1A. We see that in the strong proofreading regime (i.e., $k_f \leq \langle k_{\text{off}}\rangle$), the mutual information decreases as $N$ increases. Furthermore, weak proofreading (i.e., a fast proofreading rate) always improves the mutual information compared to the strong proofreading regime. The mutual information therefore supports the picture we saw with the SNR, that strong proofreading and large $N$ actually decrease the fidelity information transmission through the receptor.

**B. Further explanation of the resolution metric.** Another way to understand the resolution metric (Eq. [6]) is to imagine that ligands are classified based on whether their estimated affinity, $k^*_{\text{off}}$, is above or below a classification threshold $k^{\text{threshold}}_{\text{off}}$. Conventionally, $k^{\text{threshold}}_{\text{off}}$ can correspond to some minimum activating number of phosphorylated molecules output by the receptor via $\langle n\rangle(k^{\text{threshold}}_{\text{off}}, c, t) = n_{\text{activating}}$. The ability of a receptor to discriminate between two ligands is then determined by the separation between estimation distributions for each ligand, since a greater separation makes estimating $k^*_{\text{off}} > k^{\text{threshold}}_{\text{off}}$ less likely when the ligand generating $n$ has unbinding rate $k_{\text{off}} < k^{\text{threshold}}_{\text{off}}$ and vice versa. Fisher's linear discriminant is designed to measure the separation between the estimation distributions for each ligand.

The full expression for an $N$-step KPR receptor's resolution as measured by the Fisher linear discriminant metric is:

$$\eta_{\text{FLD}} = \frac{k_p t x(\delta-1)^2}{A+B} \qquad [14]$$

$$\text{where } A = \frac{(1+g)^{2+N}(1+2k_p/k_{\text{off}})(1+x)^3}{\Big(1+g(1+N+Nx)\Big)^2} - \frac{2(1+g)k_p x(1+x)(2+x+g(2+N+x+Nx))}{\Big(1+g(1+N+Nx)\Big)^2}$$

$$\text{and } B = \frac{(x+\delta)(1+g\delta)}{\delta\Big(1+g\big(\delta+N(x+\delta)\big)\Big)^2}\Bigg( (x+\delta)^2(1+g\delta)^{N+1}(2\frac{k_p}{k_{\text{off}}}+\delta) - 2k_p x\Big(x+g(1+N)x\delta+\delta(2+g(2+N)\delta)\Big)\Bigg)$$

Figures S1 (G-H) indicates where the resolution of the proofreading receptor is worse than a non-proofreading receptor; that is, in the low concentration ($y < 1$) or strong proofreading regimes. While the ratio $\eta^N_{FLD}/\eta^0_{FLD}$ (Eq. [6] of the main text) is monotonic in $g$ and independent of ligand concentration $c$, Fig. S1 H reveals a more complex dependence on these parameters when noise is taken into account. Note that the ratio $\eta^N_{\text{FLD}}/\eta^0_{\text{FLD}}$ is independent of the time at which the signal $n$ is observed, so waiting for receptor output signal to accumulate does not abrogate the effect of noise or in any other way improve resolution.

**C. Stabilized signaling state.** The variance for a 1-step KPR receptor with a stabilized signaling state is, at long times,

$$\sigma_n^2 = \frac{k_p t x}{(x+\alpha+g\alpha+gx\alpha)^3}\Bigg((x+\alpha+g\alpha+gx\alpha)^2 + 2\alpha\frac{k_p}{k_{\text{off}}}\Big(1+g^2(1+x)^2+g(2+x)\Big)\Bigg) \qquad [15]$$

The width of the output distribution is quantified by the ratio of the mean over the standard deviation. An expression for the general $N$-step stabilized proofreading receptor model is not known but the mean and variance were computed by Gillespie simulation and plotted in Fig. S2 C versus the theoretical scaling $1/g^{N/2}$ and was found to be in good agreement.

**D. Means and variances for KPR receptor with burst output.** The Master equation for the burst type KPR receptor is given by:

$$\frac{\partial P_0^m(c,k_{\text{off}},t)}{\partial t} = -k_{\text{on}}cP_0^m + k_{\text{off}}\sum_i^N P_i^m$$

$$\frac{\partial P_1^m(c,k_{\text{off}},t)}{\partial t} = k_{\text{on}}cP_0^m - (k_f+k_{\text{off}})P_1^m$$

$$\frac{\partial P_2^m(c,k_{\text{off}},t)}{\partial t} = k_f P_1^n - (k_f+k_{\text{off}})P_2^m$$

$$\vdots$$

$$\frac{\partial P_{N+1}^m(c,k_{\text{off}},t)}{\partial t} = k_f P_N^{m-\beta} - k_{\text{off}}P_{N+1}^m$$

where $\beta$ is the burst size and all other parameters are the same as the original KPR scheme. We solve for $\beta = 1$ and use the fact that this solution counts the number of burst events so that the total signal mean and variance can be scaled by $\beta$ and $\beta^2$ respectively.

The moments of the burst type KPR receptor are computed by solving the ordinary differential equations $\partial_t\langle m\rangle = \sum_{m=0}^{\infty}\sum_{i=0}^{N+1} m\partial_t P_i^m$ and $\partial_t\langle m^2\rangle = \sum_{m=0}^{\infty}\sum_{i=0}^{N+1} m^2\partial_t P_i^m$ (see Methods). The resulting mean, $\langle m\rangle$, and variance, $\sigma^2_{m,N}$, are provided for $N$ from 0 to 4 (with $\beta = 1$):

$$\langle m\rangle_N = \frac{k_{\text{off}}tx}{(1+g)^N(1+x)}$$

$$\sigma^2_{m,0} = \frac{k_{\text{off}}tx(1+x^2)}{(1+x)^3}$$

$$\sigma^2_{m,1} = \frac{k_{\text{off}}tx(1+2g+x^2+g^2(1+x)^2)}{(1+g)^3(1+x)^3}$$

$$\sigma^2_{m,2} = \frac{k_{\text{off}}tx}{(1+g)^5(1+x)^3}\Big((1+g)^3+2g^2(3+g)x+ (1+g(-1+g(3+g)))x^2\Big)$$

$$\sigma^2_{m,3} = \frac{k_{\text{off}}tx}{(1+g)^7(1+x)^3}\Big((1+g)^4+2g^2x(6+g(4+g))+ (1+g(-2+g(6+g(4+g))))x^2\Big)$$

$$\sigma^2_{m,4} = \frac{k_{\text{off}}tx}{(1+g)^9(1+x)^3}\Big(1+x^2+10g^2(1+x)^2+ 10g^3(1+x)^2+5g^4(1+x)^2+g^5(1+x)^2+g(5-3x^2)\Big)$$

where $g = k_{\text{off}}/k_f$ and $x = k_{\text{on}}c/k_{\text{off}}$.

**E. A minimal noise KPR model.** We can go one step further than the KPR model with fixed residency time described in Section 2F of the main text by making the time for *each* reaction other than ligand binding (which must be exponentially distributed if it is diffusion limited) happen in a corresponding deterministic time. In this model the probability for the receptor to move from one bound state to the next is $p_{i\rightarrow i+1} = k_f/(k_f+k_{\text{off}})$ and the probability for the ligand to unbind instead is $p_{\text{off}} = k_{\text{off}}/(k_f + k_{\text{off}})$. If the ligand unbinds, it happens in time $\tau = 1/k_{\text{off}}$. If the receptor moves to the next state then it happens in time $\tau = 1/k_f$. This model avoids all but the minimum noise directly associated with the proofreading dynamics. We show in Figure S2 that the resolution of this minimal noise model, measured by $\eta_{\text{FLD},n} = \big(\langle n(k_{\text{off},1})\rangle - \langle n(k_{\text{off},2})\rangle\big)^2/(\sigma^2_{n,1}+\sigma^2_{n,2})$ still scales inversely with the number of proofreading steps, $N$.

At high ligand concentrations, a fixed residency time does not improve resolution because in the high concentration regime there is competition for the receptor signaling state; a stochastic binding time allows for a quick succession of binding events which amplifies the output and thereby reduces shot noise (see Figure S2B).

**F. Effect of decreasing proofreading strength on the noise associated with proofreading steps.** It is of interest to ask whether the detrimental effects of increasing $N$ can be compensated by simultaneously decreasing $g$, while the HN resolution enhancement is fixed to a biologically required amount for a pair of ligands. This scenario is depicted in Figure S1 G which shows the fold change in $\eta^N_{\text{FLD}}/\eta^0_{\text{FLD}}$. The dashed line is the line of constant fold-change in HN resolution, $\eta^N_{\text{HN}}/\eta^0_{\text{HN}} = 4$. It indicates that any apparent increase in $\eta_{\text{FLD}}$ along this contour arises as a result of the accompanying exponential decrease in the proofreading strength provided by increasing $k_f$. At very large $N$ the required $g$ becomes exponentially small; effectively, the receptor is proofreading infinitely fast in this regime (see also Ref [38]).

The maximum fold-enhancement in resolution in Figure S1 G is approximately $10^{0.25}$, significantly lower than the HN metric would indicate. Along the contour of constant HN resolution, the FLD resolution enhancement $\eta^N_{\text{FLD}}/\eta^0_{\text{FLD}}$ remains small and saturates at large $N$ to:

$$\frac{\eta^N_{\text{FLD}}}{\eta^0_{\text{FLD}}} \approx \frac{\tilde{k} + 2 + 2\zeta^2 + \tilde{k}\zeta^3}{(\zeta - 1)\Big(\frac{C_1^{1/(1-\zeta)}(\tilde{k}+2)(\zeta-1)}{(1-\zeta+\log C_1)^2} + \frac{C_1^{\zeta/(1-\zeta)}(\zeta-1)\zeta^2(2+\tilde{k}\zeta)}{(1-\zeta+\zeta\log C_1)^2}\Big)}$$

where $\tilde{k} = k_{\text{off}}/k_p$, $C_1 = \eta^N_{\text{HN}}/\eta^0_{\text{HN}}$, and we have taken $x = c/K_D \ll 1$. Overall, this figure confirms the main conclusion in the main text: moderate proofreading benefits for ligand discrimination are realized only in a narrow region of $(g, N)$ space, typically for low $g$ and $N$.

**G. Ligand resolution for short signaling times.** The metric $\eta_{\text{FLD}}$ can only be investigated at short times by stochastic simulation since a closed form expression is not known for the time-dependent solution of the Master equation (Eq. [11] of the main text) with an unbound receptor as the starting condition. The mean and variance of the receptor output at short times was simulated by Gillespie algorithm and the resolution metric was computed from 5000 simulations at each point in Figure 4B of the main text.

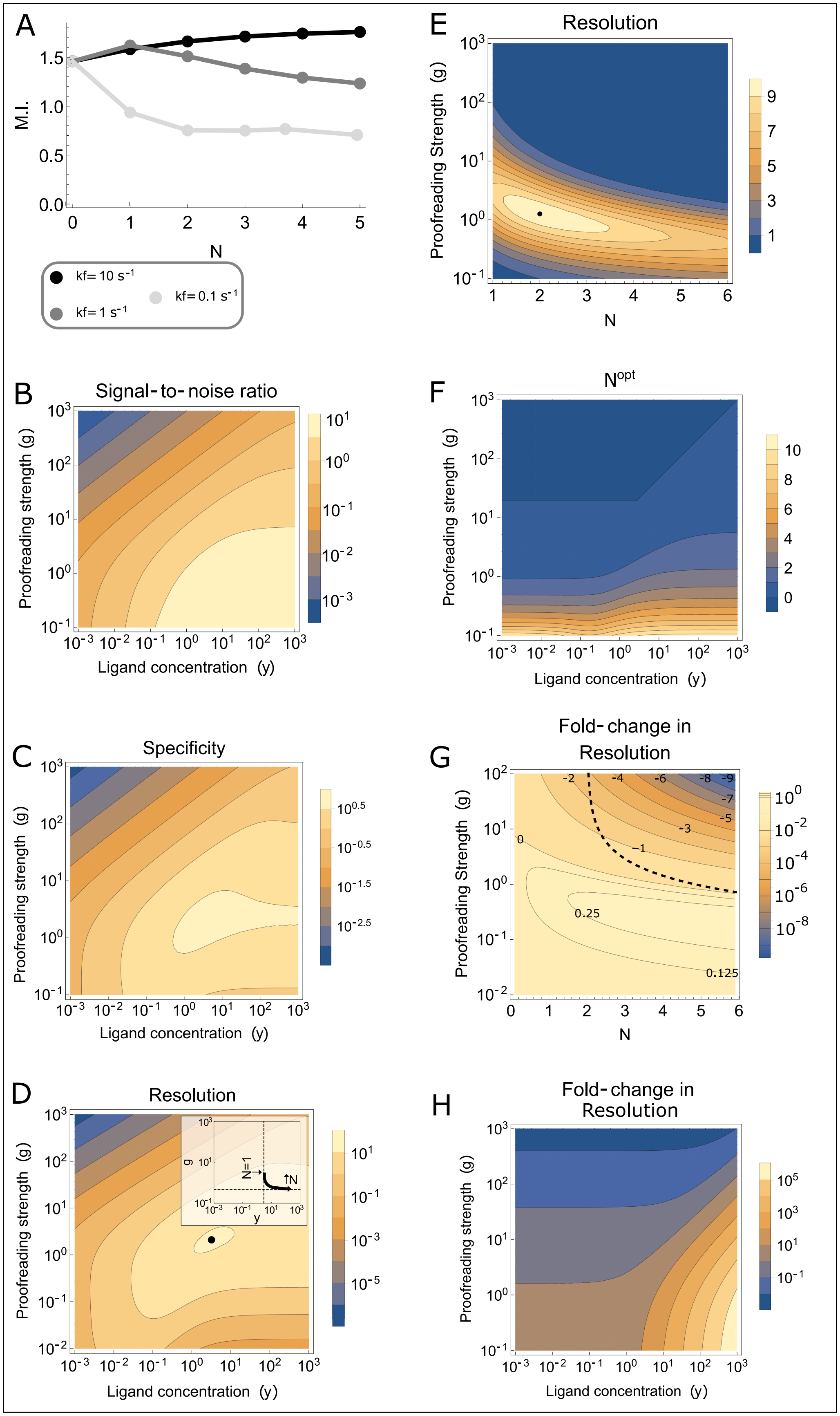


**Fig. S1.** (Caption on the following page.)

**Fig. S1.** A) The mutual information (M.I.) of the KPR receptor, $\mathcal{I}(n, k_{\text{off}}$, is computed from a log-normal prior on $k_{\text{off}}$ with unit variance and a conditional probability $P(n|k_{\text{off}}) = \mathcal{N}(\langle n \rangle_N, \sqrt{\sigma_n^2})$ for the receptor output. The M.I. decreases as the proofreading rate $k_f$ decreases (proofreading strength increases), for all choices of the number of proofreading steps, $N$. B) The signal-to-noise ratio $\langle n \rangle / \sigma_n$ for a KPR receptor as a function of the number of ligand concentration, $y$, and proofreading strength, $g = k_{\text{off}}/k_f$, with $k_{\text{on}}c/k_p = 1$, $k_f/k_p = 1$, and $k_p t = 100$. C) The specificity $k^*_{\text{off}}/\sigma_{k_{\text{off}}}$ for the estimation distribution as a function of proofreading strength, $g$, and ligand concentration, $y$, using $n = \langle n \rangle (k_{\text{off}})$ and $g = k_{\text{off}}/k_f$ for the 'true' ligand generating the output $n$. Other parameters: $k_{\text{on}}/k_p = 1M^{-1}$, $k_f/k_p = 1$, and $k_p t = 100$. D) The resolution of a KPR receptor with $N = 1$ and $\zeta = 0.1$, as measured by $\eta_{\text{FLD}}$. Resolution is maximized, for $N = 1$, at $(y_{\text{max}}, g_{\text{max}}) = (3.2, 2.1)$, indicated by black point. Inset: the location of the point of maximum resolution in the $(y, g)$-plane is plotted for $N \in [0.75, 4]$, with $\zeta = 0.1$ and $k_p t = 100$. The dashed lines show the asymptotic values of $g_{\text{max}}$ and $y_{\text{max}}$ in the large $N$ and small $N$ limits, respectively. E) The resolution of a KPR receptor in the high ligand concentration regime: $k_{\text{on}}c/k_p = 100$, $k_p t = 100$, $k_f/k_p = 1$, and $\zeta = 0.1$. The resolution has a maximum, indicated by black point, which depends on the ligand concentration but remains in the weak proofreading regime similarly to Figure 3A in main text. F) The fold-enhancement to resolution, as measured by $\eta_{\text{FLD}}$, can have a maximum at small positive values of $N$ (the number of proofreading steps) for some choices of ligand concentration $y$ and proofreading strength $g$. The optimal number of proofreading steps, $N^{opt}$, which achieves this maximization tends to increase with decreasing proofreading strength. This is unlike the results for the metric $\eta_{\text{HN}}$ where resolution monotonically increases with $N$ for all conditions. Here we have chosen the parameters $k_{\text{on}}/k_p = 1M^{-1}$, $k_f/k_p = 1$, $\zeta = 0.1$, and $k_p t = 100$. G) The fold-change in resolution, measured by $\eta^N_{\text{FLD}}/\eta^0_{\text{FLD}}$ as a function of proofreading strength $g$ and number of proofreading steps $N$ (for fixed $k_{\text{off}}/k_p = 1$ and $\zeta = 0.5$, and varying $k_f$). Contours are labeled with the $\log_{10}$ value of $\eta^N_{\text{FLD}}/\eta^0_{\text{FLD}}$. A contour of constant fold-change in HN resolution, $\eta^N_{\text{HN}}/\eta^0_{\text{HN}} = 4$, is provided for comparison (black dashed line). The concentration is the same as that of Figure 3 in the main text. $k_{\text{on}}c/k_{\text{off}} = 1$, and $k_p t = 100$. H) The fold-change in resolution by the addition of one proofreading step compared to a non-proofreading receptor, as measured by $\eta^1_{\text{FLD}}/\eta^0_{\text{FLD}}$. The orange region of parameter space indicates better resolution for a receptor with proofreading than one without proofreading. Similar results are obtained for additional proofreading steps.

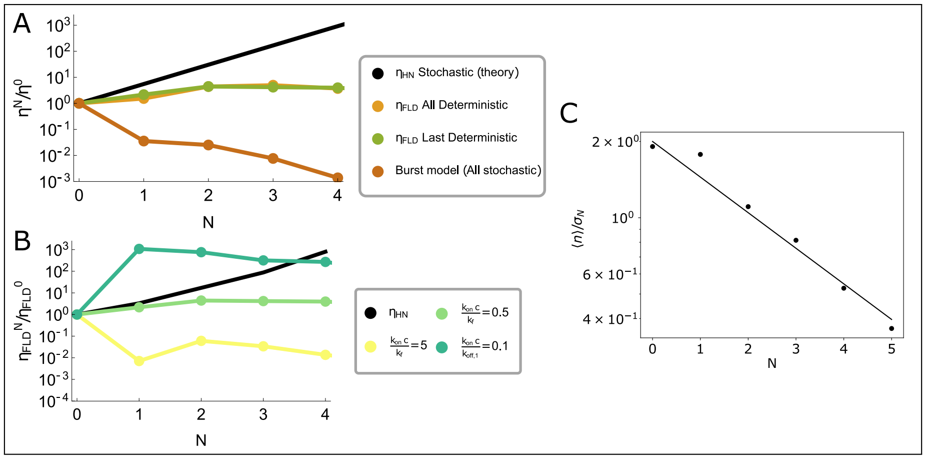


**Fig. S2.** A) The resolution of the minimal noise burst-type signal KPR scheme (with all time steps deterministic) compared to the burst models treated in the main text (last step deterministic or completely stochastic). The classical Hopfield resolution is provided for comparison; the theoretical result (Eq. [8] in the main text) and the simulated result (the ratio of the mean output for each ligand) agree very closely, as do the values of $\eta_{\text{FLD}}$ for the models with deterministic time steps. The proofreading strength is $g = k_{\text{off},1}/k_f = 10$, the burst size is 10, $k_{\text{off},1}/k_{\text{off},2} = 0.1$, $k_f/k_{\text{on}}c = 2$, and $k_{\text{on}}ct = 500$. B) The resolution enhancement of the burst + fixed residence time KPR model, measured as $\eta^N_{n,\text{FLD}}/\eta^0_{n,\text{FLD}}$, using the receptor output $\mathcal{N}(\langle n \rangle, \sigma_n^2)$ from stochastic simulation, for increasing ligand concentrations. The Hopfield resolution at $k_{\text{on}}c/k_{\text{off},1} = 0.5$ is provided (black) for comparison. The proofreading strength is $g = k_{\text{off},1}/k_f = 10$, the burst size is 10, $k_{\text{off},1}/k_{\text{off},2} = 0.1$, and $k_f t = 100$. At $N = 1$ the mean signals are equal (and therefore indistinguishable) when $k_{\text{on}}c/k_f = 10$, which is why the resolution at $N = 1$ for $k_{\text{on}}c/k_f = 5$ (close to indistinguishable) is particularly low. In general, the mean G-protein type output for $N = 1$ will be indistinguishable when $k_{\text{on}}c = k^2_{\text{off}}\zeta/k_f$. C) The ratio $\langle n \rangle/\sigma_N$ for the stabilized proofreading receptor model, as a function of $N$ the number of proofreading steps. Means and variances were computed from 300 independent Gillespie simulations (points) and the theoretical scaling $(\langle n \rangle_0/\sigma_0)^{N/2}$ was plotted for comparison (line). The agreement demonstrates that the stabilized proofreading model does not overcome the adverse effects of proofreading on the output distribution noise.
